# ciRS-7 exonic sequence is embedded in a long non-coding RNA locus

**DOI:** 10.1101/169508

**Authors:** Steven P. Barrett, Kevin R. Parker, Caroline Horn, Miguel Mata, Julia Salzman

## Abstract

ciRS-7 is an intensely studied, highly expressed and conserved circRNA. Essentially nothing is known about its biogenesis, including the location of its promoter. A prevailing assumption has been that ciRS-7 is an exceptional circRNA because it is transcribed from a locus lacking any mature linear RNA transcripts of the same sense. Our interest in the biogenesis of ciRS-7 led us to develop an algorithm to define its promoter. This approach predicted that the human ciRS-7 promoter coincides with that of the long non-coding RNA, LINC00632. We validated this prediction using multiple orthogonal experimental assays. We also used computational approaches and experimental validation to establish that ciRS-7 exonic sequence is embedded in linear transcripts that are flanked by cryptic exons in both human and mouse. Together, this experimental and computational evidence generate a new view of regulation in this locus: (a) ciRS-7 is like other circRNAs, as it is spliced into linear transcripts; (b) expression of ciRS-7 is primarily determined by the chromatin state of LINC00632 promoters; (c) transcription and splicing factors sufficient for ciRS-7 biogenesis are expressed in cells that lack detectable ciRS-7 expression. These findings have significant implications for the study of the regulation and function of ciRS-7, and the analytic framework we developed to jointly analyze RNA-seq and ChIP-seq data reveal the potential for genome-wide discovery of important biological regulation missed in current reference annotations.

**Author Summary:** circRNAs were recently discovered to be a significant product of ‘host’ gene expression programs but little is known about their transcriptional regulation. Here, we have studied the expression of a well-known circRNA named ciRS-7. ciRS-7 has an unusual function for a circRNA; it is believed to be a miRNA sponge. Previously, ciRS-7 was thought to be transcribed from a locus lacking any mature linear isoforms, unlike all other circular RNAs known to be expressed in human cells. However, we have found this to be false; using a combination of bioinformatic and experimental genetic approaches, in both human and mouse, we discovered that linear transcripts containing the ciRS-7 exonic sequence, linking it to upstream genes. This suggests the potential for additional functional roles of this important locus and provides critical information to begin study on the biogenesis of ciRS-7.

## Introduction

Until recently, the expression of circRNA was almost completely uncharacterized, with a few important exceptions [1–4]. It is now appreciated that circRNAs are a ubiquitous feature of eukaryotic gene expression [1,3,5,6]. While many functions have been posited for circRNAs, few have been supported with experimental evidence. ciRS-7, one of the most highly expressed and most intensely studied circRNAs, is an exception to this rule, where recent work has shown it functions as a miRNA sponge [6,7]. The sequence of ciRS-7 is highly repetitive with over 70 repeated miR-7 seed sequences in humans, most of which are conserved across eutherian mammals, and its expression is highly variable across tissues [6–8]. ciRS-7 also exhibits increasing expression in neuronal differentiation models *in vitro [9]*. In zebrafish, which do not have an endogenous copy of ciRS-7 but do express miR-7, ectopic expression of the ciRS-7 sequence results in a defect in midbrain development [6]. And a recent ciRS-7 knock-out mouse exhibited neuronal defects, including impaired sensorimotor gating and dysfunctional synaptic transmission [10]. In spite of these functional findings, a model for the biogenesis and regulation of ciRS-7 is lacking, and key questions remain: What is the primary transcript that is processed to produce ciRS-7? Where in the genome is the promoter for this transcript? Are there any other spliced transcripts, circular or linear, generated from this locus?

Correlative and mini-gene analyses have suggested that biogenesis of some circRNAs is regulated by intronic sequence flanking the circularized exon [11–13]. However, the immediate flanking sequence does not appear to control the biogenesis of ciRS-7; inserting 1 kb of the endogenous sequence flanking the ciRS-7 exon into a plasmid driven by a CMV promoter was not sufficient to produce ciRS-7 [7], implying that additional sequence is necessary for circularization. Identifying this additional sequence is an especially difficult problem in the case of ciRS-7, as the intron upstream of the circularized exon has not been described due to the lack of an annotated promoter, and unlike every other known human circular RNA, ciRS-7 is not thought to be included in a mature linear transcript, obscuring possible transcriptional start sites that would be shared with these isoforms.

Identifying the promoter for ciRS-7 has broad implications but is non-trivial: unlike for linear RNAs where techniques like 5’ RACE can determine the transcription start site (TSS), no such approach can be used for ciRS-7 or any other circRNA. To overcome this problem and to identify the TSS of ciRS-7, we designed a new statistical method that entailed integrative analysis of chromatin modifications measure by ChIP and the RNA expression levels of ciRS-7 to identifying its promoter. This is a general analytic framework that could be applied to any transcript, but we chose to focus on ciRS-7 because of the biological significance described above and because it has stood out as the only case of a human circRNA with no known linear counterpart.

Our analysis led us to discover that the promoters of a nearby uncharacterized locus currently annotated as a long non-coding RNA (LINC00632) were responsible for driving ciRS-7 expression. In contrast to current thinking in the field, we also discovered that, in both human and mouse, the ciRS-7 exon is embedded in novel linear transcripts that include cryptic exons both up and downstream of the ciRS-7 sequence. In humans, these linear transcripts include exons overlapping with LINC00632, and the subcellular localization of transcripts from LINC00632 vary depending on the presence of the ciRS-7 sequence.

Together, these results support (post-)transcriptional coupling between a long-noncoding RNA and ciRS-7 and raise important functional questions about this locus. Our results also represent the first steps toward pinpointing the mechanisms underlying the regulation and biogenesis of ciRS-7.

## Results

### Computational methods predict the ciRS-7 promoter region

As a first step to identify the ciRS-7 promoter, we examined available chromatin immunoprecipitation sequencing (ChIP-seq) data from HeLa, HEK293, and *in vitro* differentiated neuronal cells, which exhibit a range of ciRS-7 expression, from very low (or nonexistent) in HeLa cells to very high in neurons [7–9].

We investigated RNA Polymerase II binding as well as the histone modifications H3K4me3 and H3K27ac, which are enriched in active promoters [14,15]. In addition, we examined H3K4me1, which is enriched in enhancers, and H3K27me3, a repressive mark enriched in silenced loci [16]. The only peaks called by MACS2, a widely used peak-calling algorithm [17], in H3K4me3, H3K27ac or RNA Polymerase II were in HEK293 and *in vitro* differentiated neuronal cells and coincided with the transcriptional start positions of LINC00632 isoforms, the nearest annotated transcript upstream of ciRS-7 (Fig 1A; S1 Fig). Conversely, the repressive mark H3K27me3 was visibly enriched throughout the LINC00632 and ciRS-7 locus in HeLa cells, consistent with their lack of ciRS-7 expression, with a H3K27me3 peak called at a LINC00632 promoter (S1 Fig). This visual inspection generated the hypothesis that the LINC00632 and ciRS-7 promoters coincide, which we went on to quantitatively test.

**Fig 1.**
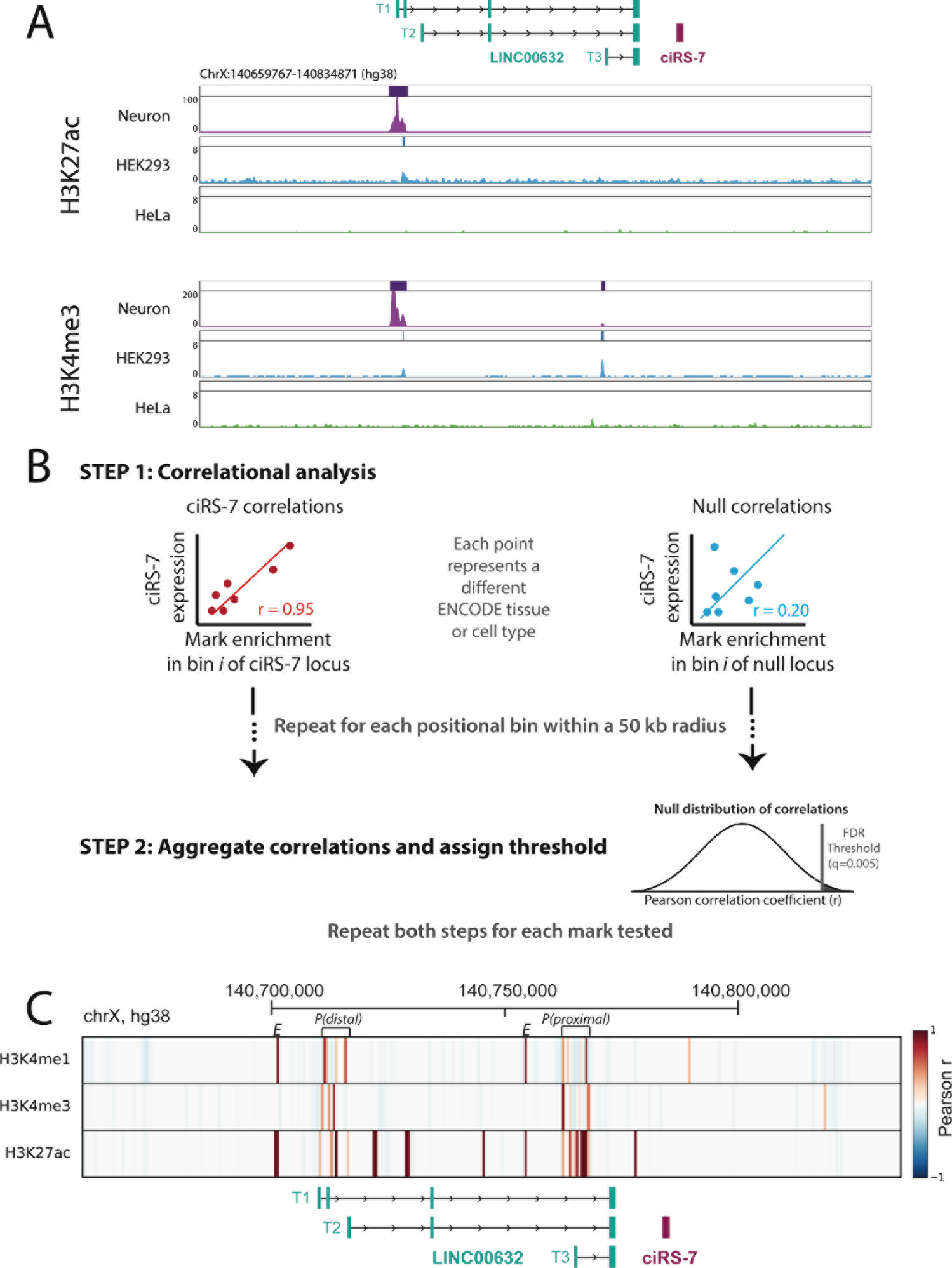
Computational and statistical analysis predict ciRS-7 shares a promoter with annotated LINC isoforms. **(A)** H3K27ac and H3K4me3 peaks across the ciRS-7 locus and nearby genomic region in *in vitro*-differentiated neurons, HEK293, and HeLa cells. **(B)** Schematic depicting analysis correlating chromatin mark enrichment near the ciRS-7 locus with ciRS-7 expression in RNA seq data. To estimate a false discovery rate, null correlations are computed using chIP-enrichment at disparate regions of the genome with no relationship to ciRS-7 (ACTB, FOXO4, and HOTAIR). Then, using these null correlations, a null distribution is created from which a false discovery rate can be estimated. **(C)** Heatmap correlation (Pearson r) between strand-specific ciRS-7 expression and enrichment of histone marks across the ciRS-7 locus and surrounding genomic region (spanning from 50 kb upstream of LINC00632 and 50 kb downstream of ciRS-7). Correlations are plotted in 500 nucleotide bins, and a depiction of annotated genes is shown below. Our FDR threshold (q<0.005) is satisfied for correlations greater than 0.35 for H3K4me1 (top 6 correlated bins), 0.45 for H3K4me3 (top 4 correlated bins), and 0.75 for H3K27ac (top 14 correlated bins) (See File S5). Putative promoters regions are annotated with brackets and the letter ‘P’. Putative enhancers are marked with the letter ‘E’

If ciRS-7 shares its promoter with LINC00632, activating chromatin marks at the LINC00632 promoter and ciRS-7 expression should be positively correlated. To test this prediction, we analyzed matched ChIP-seq and RNA-seq data from 34 ENCODE tissues and cell types. Specifically, we separated the ~175 kb genomic region spanning 50 kb upstream of the LINC00632 annotation to 50 kb downstream of ciRS-7into 500 bp bins, and we computed the Pearson correlation between ciRS-7 expression and the enrichment of each of three activating marks (H3K4me3, H3K4me1, and H3K27ac) in each bin (Fig 1B). Because the null distribution of the Pearson correlation requires assumptions that do not hold for our data, we computed an empirical null distribution: for each activating mark, we computed the Pearson correlation between ciRS-7 expression and its enrichment per bin 50 kb up- and downstream of genes that should have no relationship to expression or chromatin modifications in the ciRS-7 locus: ACTB, HOTAIR, and FOXO4. This empirical null distribution was used to estimate the FDR for the correlation coefficients at each bin/mark pair (S2 Fig, Methods).

The Pearson correlation between H3K4me3 marks and ciRS-7 expression were highest and statistically significant (q<0.005) at the two promoters of LINC00632 ( Fig 1C, marked as ‘P(Distal)’ and ‘P(Proximal)’; S3 Fig) and H3K27ac marks in these regions were also high and statistically significant (q<0.005). Coincident H3K4me1 and H3K27ac marks, marks of active enhancers [18], found in this loci were also significantly correlated with ciRS-7 expression (Fig 1C, marked as ‘E’). No significant signal at regions more proximal to the ciRS-7 exon or upstream of annotated LINC00632 isoforms were observed, providing further statistical support that ciRS-7 expression is driven from these promoters.

### Orthogonal experimental tests validate that ciRS-7 shares a promoter with LINC00632

To determine whether specific activation of the LINC00632 promoters was sufficient to drive ciRS-7 expression, we used three experimental tests. First, we used the promoter-activating CRISPRa system in HeLa cells which express low or undetectable levels of ciRS-7[7,19] to test if ciRS-7 expression could be driven by these promoters. We designed single guide RNAs (sgRNAs) to target the two promoter regions highlighted in Fig 1C, just upstream of T1 and T3, identified as the putative promoter regions by computational analysis (See S1 File for sgRNA sequences). Targeting of the CRISPRa system to either region resulted in induction of specific LINC00632 isoforms, and activating either of these promoters induced robust expression of ciRS-7 (Fig 2A), with a ΔCt compared to ACTB of ~10 (S4 Fig), although at ~2-3 orders of magnitude less than in the highly expressing HEK293 cells.

**Fig 2.**
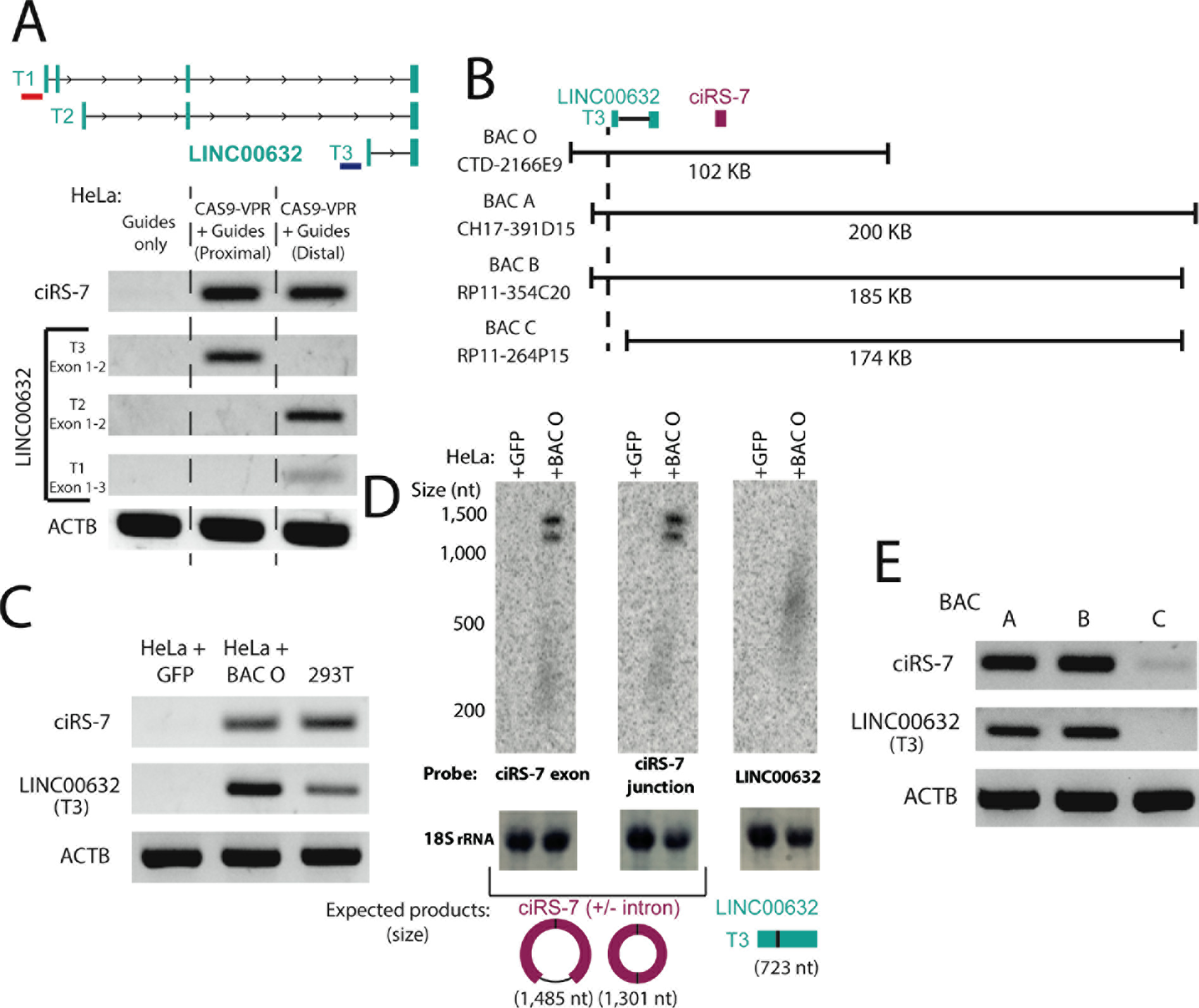
Experimental approaches confirm the identity of the ciRS-7 promoter. **(A)** RT-PCR for ciRS-7 and LINC00632 transcripts in HeLa (+/− Cas9-VPR activation) with guide RNAs targeting distal and proximal promoters identified in Fig 1C. The approximate targeted locations of the proximal and distal guide RNAs are shown with a blue and red bar, respectively, in the diagram shown above the gel. **(B)** Schematic of BAC inserts with respect to the ciRS-7 and LINC00632 genes. **(C)** RT-PCR and **(D)** Northern blot for ciRS-7 and LINC00632 transcripts generated after BAC O transfection. The appearance of two bands in the ciRS-7 Northern blots are due to alternative splicing of an intron contained within the ciRS-7 exonic sequence. **(E)** RT-PCR for ciRS-7 and LiNc00632 transcripts from HeLa cells transfected with BACs A-C.

To test the hypothesis that HeLa cells are competent to express ciRS-7 at the nearest identified promoter if it is free from repressive chromatin marks, we transfected a Bacterial Artificial Chromosome (BAC O, Fig 2B), containing a genomic fragment starting upstream of the proximal LINC00632 promoter and ending ~50 kb downstream of ciRS-7, into HeLa cells. We detected significant expression of ciRS-7 and LINC00632 from this BAC by both RT-PCR and Northern blot after one day of transfection. As an aside, this experiment supports the model that the lack of LINC00632 and ciRS-7 expression in HeLa cells is due to chromatin modification of the locus, rather than the absence of necessary trans-acting factors and shows that all (post-)transcriptional machinery required for ciRS-7 expression are present in HeLa cells (Fig 2C,D; S5 Fig).

As an orthogonal test that the proximal LINC00632 promoter, and no further downstream promoter drives ciRS-7 expression, we transfected three other BACs (A-C) each containing the exonic sequence of ciRS-7, but differing in their inclusion of the proximal endogenous promoter of LINC00632 (Fig 2B,E). Because BAC O transfects more efficiently than BACs A-C, likely due to its relatively small size, we excluded it in this comparison (S6 Fig). Expression of LINC00632 isoforms and ciRS-7 were correlated and highly dependent on inclusion of the LINC00632 promoter, further supporting the hypothesis that ciRS-7 and LINC00632 isoforms share the same promoters (Fig 2E, S7A Fig).

As a final test that the dominant promoter of ciRS-7 is located in the LINC00632 promoter, we created a genomic deletion in HEK293T cells, which express ciRS-7 at high levels [8], that encompasses the predicted positions of both putative promoters and encompass the guide RNAs used for CRISPRa (hg38 coordinates, chrX:140,709,590-140,749,836) (see diagram in S7B Fig). We obtained one homozygous clone. ciRS-7 decreased by approximately one thousand-fold (S7B Fig), but was not completely abolished, reflecting residual low-level promoter activity in the LINC00632 locus.

### ciRS-7 is an exon embedded in mature human and mouse linear RNA transcripts

In parallel with our computational approach to identify the promoter of ciRS-7, we tested whether gaps in the reference annotation of human exons might have missed upstream and or downstream exons spliced to and/or from ciRS-7. Indeed, spliceosomal circRNAs contain both 5’ and 3’ splice sites flanking their exons, and in almost all known cases, circRNA exons are embedded in linear transcripts spliced to and from downstream and upstream exons [1,3].

We developed an analysis of RNA-seq reads capable of detecting splicing outside of annotated exonic sequences, as most algorithms that do not use annotations have high false positive and negative rates [20]. We took a transparent, simple two-step approach in an effort to establish existence of exons spliced to and from ciRS-7: In step 1, all reads are aligned to the full genome and transcriptome; in step 2, unaligned reads are broken into two pseudo paired-end reads and aligned to a 100 kb radius of ciRS-7 on chromosome X (Fig 3, upper; see Methods for details). Rather than attempting to pinpoint a genomic breakpoint, this algorithm reports all reads whose pseudo-paired ends map to the reference genome (File S2). Differences in alignment positions of the two pseudo paired-end reads are used to infer the existence of splicing events. As a positive control for this algorithm, we omitted the ciRS-7 junction from our transcriptome index and attempted to (re-)discover this junction *de novo* and other un-annotated splicing events.

**Fig 3.**
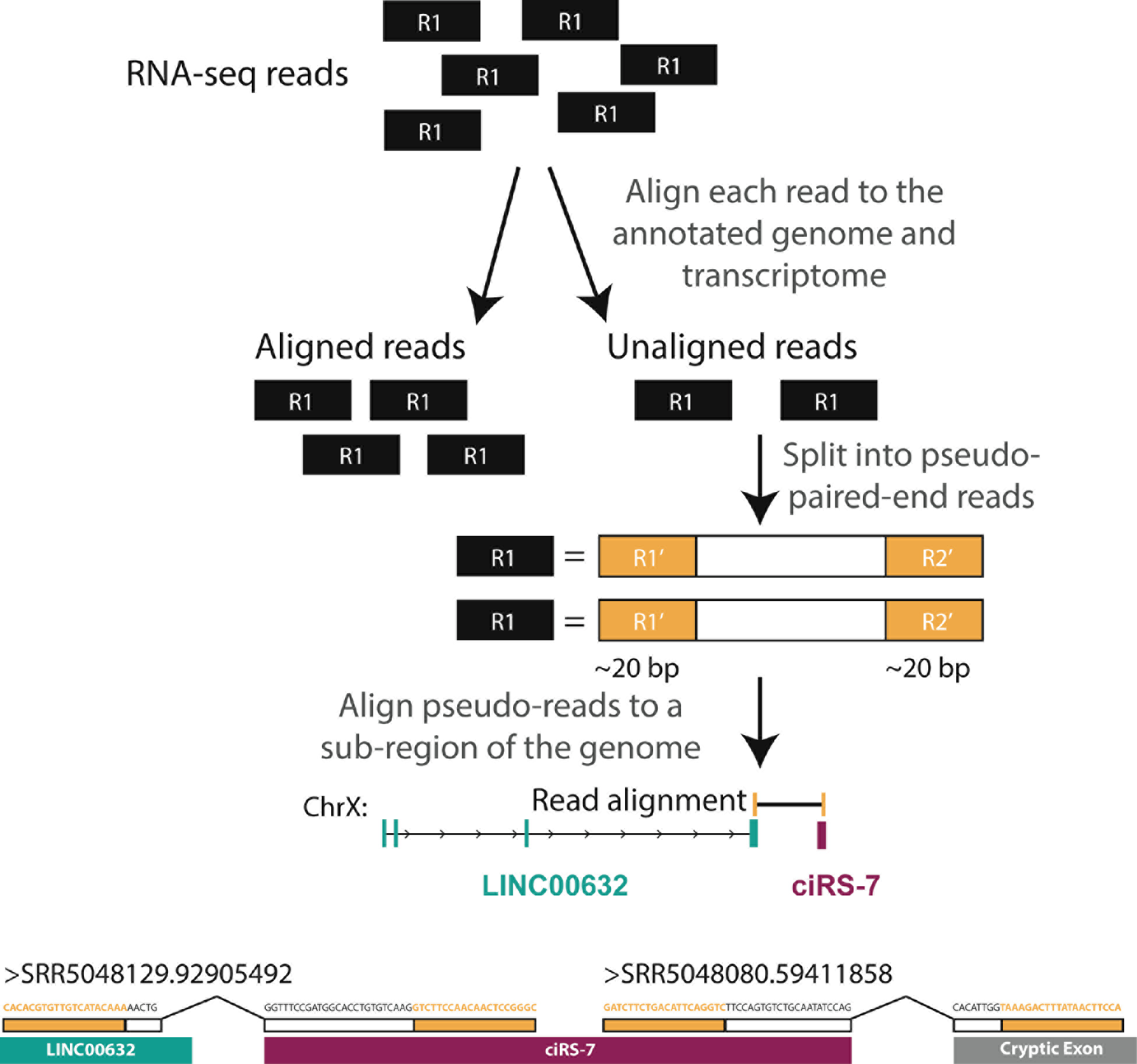
(Upper) Schematic outlining RNA-seq split read mapping approach. (Lower) Unannotated linear splicing is predicted up- and downstream of the ciRS-7 exon. Reads supporting these junctions are shown above the locus. Note, features are not drawn to scale.

We applied this algorithm to an H1 hESC RNA-seq dataset (Methods). 58 total reads representing possible novel junctions near the ciRS-7 locus were identified, 55 of which could be explained by the ciRS-7 circle junction. Of the three remaining reads, one represented a novel junction between a cryptic donor site in the final exon of LINC00632 and the ciRS-7 acceptor and a second predicted splicing between the 3’ end of ciRS-7 to a cryptic downstream exon lacking any annotation (Fig 3, lower); a final read mapped internally to ciRS-7.

To test these novel predicted splicing events, we performed RT-PCR using primers in HEK293T (Fig 4A,B; S8 Fig), a cell line known to express ciRS-7 [8]. Direct Sanger sequencing of resulting products validated our predictions, and included two cryptic 5’ splice sites in the final exon of LINC00632 paired with the annotated acceptor of ciRS-7 (S9 Fig). RT-PCR for the downstream exon yielded two splice isoforms, one of which was predicted computationally and the other using an acceptor ~1 kb upstream (Fig 4C, S9 Fig, S10 Fig). qPCR for three variants: LINC00632, the LINC00632-ciRS-7 transcript, and ciRS-7 in HEK293T showed that ciRS-7 was ~250-fold more abundant than the LINC00632-ciRS-7 transcript and about ~50-fold more abundant than LINC00632 (S11 Fig). LINC00632-ciRS-7 and LINC00632 transcripts were also RNase R sensitive, as expected for linear transcripts (S12 Fig).

**Fig 4.**
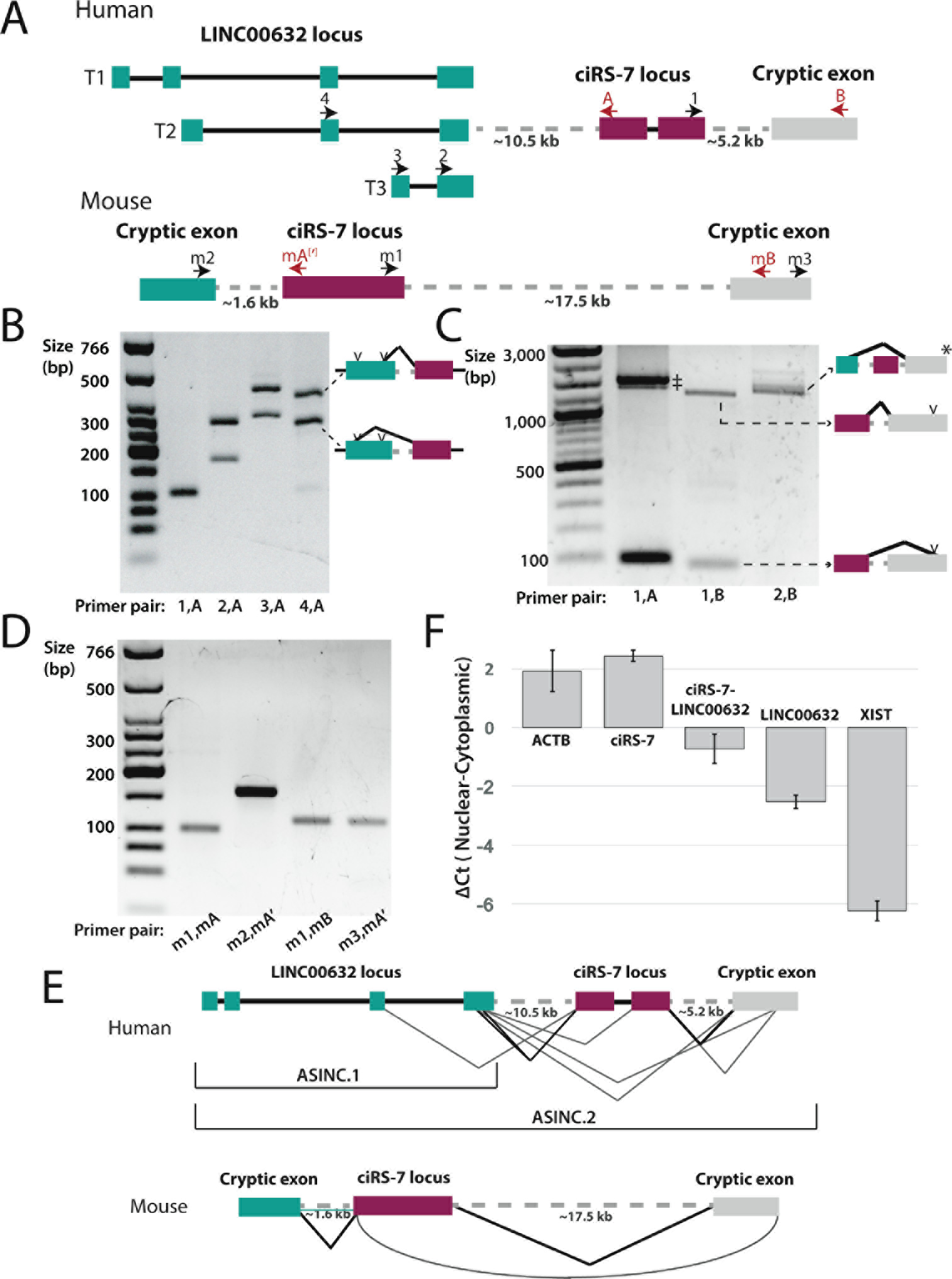
ciRS-7 exonic sequence is included in linear transcripts. **(A)** Schematic of the locus including PCR primers used in this study. Dotted lines indicate approximate positions of newly discovered introns. Figure is not drawn to scale. **(B)** RT-PCR of circular and linear ciRS-7 splice products from HEK293T. (1,A): control PCR for ciRS-7; other lanes: LINC00632 exons spliced to the ciRS-7 exonic sequence: the two bands in each of these lanes represent the products formed when the two possible splice sites in the final exon of LINC00632 are used (see diagram on the right of gel). **(C)** PCR of spliced products that include cryptic exons downstream of ciRS-7. (‡) represents rolling circle ciRS-7 PCR products with and without intron retention. (*) Other products were also identified (see S7 Fig). **(D)** RT-PCR of circular and linear ciRS-7 splice products from mouse brain RNA. mA and mA’ bind to approximately the same region but have slightly different sequence (see S1 File). **(E)** Examples of novel splicing observed in the human and mouse ASINC loci. Curved line in mouse indicates a backsplice. **(F)** qPCR quantification of nuclear-cytoplasmic fractionated RNA from HEK293T. Error bars represent the standard deviation of biological replicates.

In an effort to test transcriptional co-regulation between LINC00632 and LINC00632-ciRS-7, we profiled 136 ENCODE cell lines and tissues to quantify both (a) the total expression of ciRS-7 sequence compared to LINC00632 and (b) the splice variants ciRS-7 and ciRS-7-LINC00632 (S13 fig). This analysis revealed that (a) ciRS-7 is more highly expressed in muscle and fat tissues than in the brain (based on TPM values); (b) the expression of LINC00632-ciRS-7 splicing versus ciRS-7 expression is tissue-specifically regulated; and (c) while relative expression levels of LINC00632 and ciRS-7 have a dynamic range across several orders of magnitude, their expression was highly correlated across all cell lines and tissues we analyzed (Pearson r=0.57, Spearman r=0.41, both p-vals≪10e-6). Together, this analysis suggests both transcriptional coupling and differential regulation of splicing between LINC00632 and ciRS-7.

Many features of ciRS-7 expression are conserved in mammals [6]. We hypothesized that its embedding in linear transcripts was similarly conserved, despite the current thinking that ciRS-7 lacks a mature linear transcript in mammals [6–8,10,21]. To test this hypothesis, we applied the the same analytic approach used above in human cells to mouse (Methods). It also predicted the existence of novel cryptic exons flanking ciRS-7, variants that were confirmed by RT-PCR in mouse brain (Fig 4D,E). In addition, it predicted a new circRNA resulting from back-splicing of a cryptic exon 15 kb downstream of ciRS-7 to its annotated acceptor, which we validated by PCR and sequencing (Fig 4D,E). qPCR for the linear junction between the novel upstream exon and ciRS-7 showed this isoform was RNase R sensitive, evidence of it being linear, and ~250 fold less abundant than ciRS-7 (S14 Fig). In this experiment, ciRS-7 was also strongly sensitive to RNase R, as has been reported by others [3]. While exonic sequences flanking ciRS-7 in linear transcripts have no detectable primary sequence homology between human and mouse, such conservation is not necessarily expected for long non-coding RNAs [22].

Our RNA-seq analysis focused on establishing the existence of cryptic exons spliced to and from the ciRS-7 exon, rather than complete annotation of all existing transcripts. We sought to estimate transcript diversity in this locus by exploratory RT-PCR in human cells. This work uncovered isoforms that splice directly from LINC00632 to cryptic exons downstream of ciRS-7, including skipping of the ciRS-7 sequence and direct splicing into the downstream internal exon of ciRS-7 (Fig 4E, S10 Fig). We attempted several PCRs not guided by the RNA-seq analysis described above; in general, these PCR reactions were negative, evidence against a model of pervasive noisy splicing in the locus, and evidence that we had identified the dominant transcripts expressed from the LINC00632 locus.

### ciRS-7 sequence has potential circRNA-independent regulatory effects

To test for potential differential regulation of ciRS-7, LINC00632, and LINC00632-ciRS-7 linear transcripts, we profiled their subcellular localization by fractionating nuclear and cytoplasmic RNA from HEK293T. Using XIST and ACTB as controls for enrichment of nuclear and cytoplasmic fractions, respectively, qPCR demonstrated that, relative to ciRS-7, LINC00632 and LINC00632-ciRS-7 were enriched in the nucleus with increasing degrees (~9 and 25-fold respectively vs. ciRS-7), suggesting that the ciRS-7 sequence impacts the steady-state localization of transcripts containing it (Fig 4F).

One hypothesis generated by this work is that the expression of ciRS-7 is directly or indirectly tied to the expression of other transcripts in this locus. The recent study that knocked out the ciRS-7 sequence in mouse allows us to begin to test this hypothesis [10].

While we have not experimentally validated the transcriptional start of the ciRS-7 pre-mRNA in mouse, there are many similarities to the human locus: the nearest upstream gene is an uncharacterized lincRNA (C230004F18Rik) and the nearest H3K4me3 (a mark of promoter activity) and RNA Polymerase II peaks to ciRS-7 occur at the transcriptional start of this lincRNA (Fig 5A). In addition, transcription of this locus appears to begin at the C230004F18Rik promoter, proceeding continuously past the ciRS-7 locus (Fig 5A). Taken together, these data suggest that the transcriptional start of the ciRS-7 pre-mRNA in mouse occurs at an upstream lincRNA, namely C230004F18Rik, as it does in human. This raises the question: is the abundance of C230004F18Rik affected by the presence or absence of the ciRS-7 sequence?

**Fig 5.**
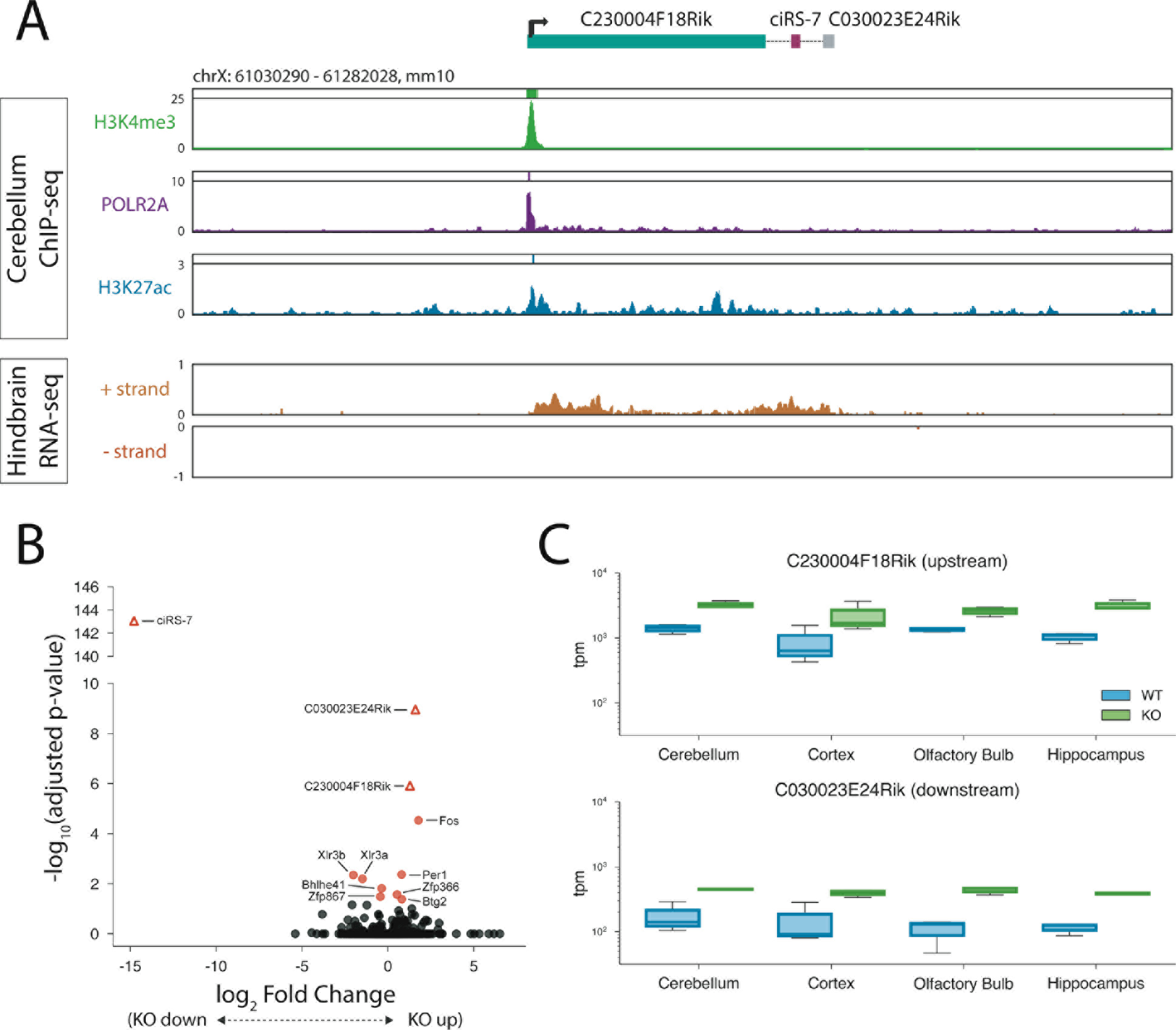
Up- and downstream transcripts are upregulated in ciRS-7 KO mouse. **(A)** ChIP-seq and RNA-seq tracks for mouse cerebellum and hindbrain in the genomic region surrounding the ciRS-7 locus. **(B)** Volcano plots of log fold-change ciRS-7 KO vs WT collapsed across four brain regions. **(C)** Box plots depicting tpm of C230004F18Rik (top) and C030023E24Rik (bottom) in WT and ciRS-7 KO mice across four brain regions.

To test this, we re-analyzed data from mouse ciRS-7 knockout experiment to determine if deletion of the ciRS-7 sequence resulted in differential expression of C230004F18Rik. We discovered that, when collapsing across all brain regions profiled, among differentially expressed genes that are statistically significant, C230004F18Rik is the third-most upregulated gene in the ciRS-7 KO vs. WT (Fig 5B,C; 2.44 fold induction, p-adjusted 1.21e-6). The most upregulated gene is Fos (3.44 fold, p-adjusted 2.95e-5), and the second-most upregulated gene, C030023E24Rik (3.03 fold, p-adjusted 1.10e-9), is an uncharacterized transcript located roughly 5.5 kb downstream of ciRS-7 and encompassed by the transcriptional read-buildup (Fig 5A-C).

In the hippocampus, C230004F18Rik was the most significantly changed gene after ciRS-7 by p-value, with 3.11 fold higher expression (p-adjusted 2.51e-13), followed by C030023E24Rik (3.4 fold higher, p-adjusted 6.29e-13) (S15 Fig). And C230004F18Rik was significantly upregulated in all tissues examined except the cortex (S15 Fig) (the lack of significant differential expression was presumably due to higher variability between replicates than in other tissues, see Fig 5C). This consistent and large effect of ciRS-7 knock-out on C230004F18Rik is consistent with a direct regulatory impact of the ciRS-7 sequence on C230004F18Rik abundance.

## Discussion

The embedding of ciRS-7 sequence in cryptic exons changes the view of the exceptionality of ciRS-7 as circRNA lacking a linear host transcript: its transcriptional regulation is similar to other circRNAs that are embedded in linear ‘host’ RNAs. In human, we have computationally and experimentally demonstrated that LINC00632 and ciRS-7 share the same promoters, and have strong support for a similar model in mouse from: (a) RNA-seq analysis showing strong expression effects on flanking linear Riken transcripts from ciRS-7 KO; (b) RNA-seq and PCR data linking ciRS-7 to cryptic up- and downstream exons; and (c) continuous RNA-seq signal across the locus starting from the upstream gene C230004F18Rik, through ciRS-7, and into the downstream transcript C030023E24Rik. Together, these data support the model that C230004F18Rik, ciRS-7 and C030023E24Rik represent a single transcriptional locus.

Because of the syntenic hosting of ciRS-7 in linear transcripts, and to simplify nomenclature, we propose renaming the uncharacterized gene LINC00632 (respectively Riken transcripts) to **<u>A</u>**lternatively **<u>S</u>**pliced **<u>IN</u>**to **<u>C</u>**iRS-7 (ASINC, respectively Asinc in mouse); we call the linear variants of these non-coding RNAs lacking ciRS-7 sequence ASINC.1 and those containing it ASINC.2.

The transcriptional and splicing machinery necessary for ciRS-7 expression is likely not brain specific, and rather is general: BACs containing a fragment of the LINC00632 locus that includes its promoter can express ciRS-7 when introduced to HeLa cells (which have little to no endogenous expression of ciRS-7). ciRS-7 expression may be primarily regulated at the level of chromatin modification of the locus, either at the newly-discovered promoters or putative enhancers.

Our work has implications for the assigned functions of ciRS-7. Despite intensive study, the promoter and mechanisms regulating ciRS-7 expression have remained mysterious. To date, experiments studying the function of the ciRS-7 sequence have made what was a reasonable simplifying assumption that the only transcripts containing the ciRS-7 sequence are circular [8,10]. However, interpretation of past and future studies assigning function to the ciRS-7 sequence must be made in light of its origination from and potential functions in linear transcripts. This includes a recent study of a mouse model where the ciRS-7 exon was deleted [10]. Our analysis of the ciRS-7 locus shows that knock-out of this exon results in an upregulation of both the upstream and downstream Riken transcripts, C230004F18Rik and C030023E24Rik. The magnitude and significance of these effects compared to other gene expression changes suggests it was a direct effect of the ciRS-7 knockout, and raises the possibility that some functions assigned to the knock-out animal could be due to increases in expression of the Riken transcripts.

There are multiple models that could explain the increased expression of the up-and downstream-Riken transcripts ciRS-7-null mice. If ciRS-7 transcription originates from a promoter shared with C230004F18Rik, as is the case for human transcription, then ciRS-7 would originate from the same linear pre-spliced transcript as the Riken transcripts. This raises the possibility that generating linear transcripts containing the ciRS-7 sequence destabilizes them or that the act of circRNA biogenesis itself leads to destabilization of the residual linear transcript. These explanations would predict upregulation of the Riken transcripts when the ciRS-7 exon is deleted. Another model is that ciRS-7 could directly or indirectly affect expression of the Riken transcripts through regulatory networks. For example, ciRS-7 may sequester or compete for splicing factors (e.g., nuclear Ago2 [23]), or ciRS-7 expression may affect the expression of other genes that are involved in direct regulation of the Riken transcripts.

The view that ciRS-7 directly regulates its host transcript is supported by our finding that ASINC transcripts are differentially localized in the cell depending on their inclusion of the ciRS-7 sequence. This suggests active regulation of or by the transcripts potentially through factors that bind the sequence in ciRS-7, and generates the hypothesis that ciRS-7-containing linear ASINC.2 has different functions in the nucleus than the ciRS-7 circRNA in the cytoplasm. Indeed, the cytoplasmic ciRS-7 circles have been shown to function by sequestering mir-7[6,7], a mechanism that unlikely to be employed by nuclear ASINC.2. Further study of transcripts in the entirety of the locus and their regulation may reveal new functions for ciRS-7 and this locus as a whole.

This work also lays a foundation for the field to begin to dissect transcriptional regulation and biogenesis of the ciRS-7 locus. Regulation of alternative splice variants in the ASINC locus, transcription factor binding patterns, and three-dimensional interactions between the promoters and putative enhancers we identified can now be analyzed across different cell types and throughout development. In addition, the analysis presented here could be generalized to other genes whose promoters are not well-annotated. For example, another well-known circRNA with no annotated promoter is derived from the Sry gene in mouse, which is circularized in mature adult testes but expresses an unspliced mRNA in the developing genital ridge that governs sex determination [2]. It has been hypothesized, though remains untested, that the promoter for the Sry circRNA uses a separate promoter from the linear mRNA[24]. Other highly-expressed circRNAs in human are also derived from the first exon of an annotated mRNA [25]. Given our results, we predict that the promoters of these circular RNAs may have been misidentified and/or misannotated in the genome and may be associated with cryptic up- or downstream splice junctions. Future analyses that comprehensively characterize transcription and splicing at individual loci will be required to fully understand the regulatory mechanisms underlying circRNA biogenesis.

## Methods

### RNA-seq analysis for detection of novel splice isoforms

Raw RNA-seq reads from SRR5048080 (human) and SRR1785046 (mouse) were downloaded from the SRA. Reads were mapped and analyzed using KNIFE [27]. Reads failing to align (“unaligned”) by KNIFE were used in a simple custom mapping approach to identify novel splicing events within a single read: single reads were split into pseudo-pairs (k-mers) by taking the first 20 mer in the unaligned read and the remaining k-mer defined by the start position O (30 for mouse and 50 for human because of differing input read lengths) and the minimum of 0+20 and the remaining read length after trimming (see supplemental perl script in S2 File). Only reads where each pseudo-pair > 18 nt were used for analysis. These pseudo-paired end reads, which actually came from the same single read were then realigned separately as pairs using bowtie2 to a custom index made by the following sequences:

>mm10_knownGene_uc012hid.1 range=chrX:61083246-61285558 5’pad=0 3’pad=0 strand=-repeatMasking=none
>hg38_knownGene_uc004fbf.2 range=chrX:140683176-140884660 5’pad=0 3’pad=0 strand=+ repeatMasking=none

Pairs of pseudo-reads mapping on the same strand were identified and used for further analysis. While pseudo-alignments are used in some RNA-seq analysis approaches, few algorithms are capable of achieving precise de novo transcript recovery due to their high false positive rates and unknown false negative rates. This approach differs from other algorithms because it (a) does not use a seed and extend approach; and (b) reads are aligned to a 100 kb radius of ciRS-7 rather than the reference genome, in principle, preventing misalignment of true ASINC reads to other loci. The complete analysis, the R script used along with output is provided in S2 File. Although our analysis is likely to include other novel splicing in the locus, we focused our attention on queries for reads that supported a splice from an un-annotated location upstream of the acceptor in ciRS-7 and from the donor in ciRS-7 to an un-annotated downstream exon using criteria on the position of discordant pseudo-paired end mappings (see R script). An example of the reads that supported these events were (in human: SRR5048129.92905492_2774860 (upstream exon); SRR5048080.59411858_2626289 (downstream exon) and in mouse: SRR1785046.9826290 (upstream exon); @SRR1785046.13497132 HWI-ST1148:158:C3UJCACXX:2:2112:5228:93046 length=50 (downstream exon backsplicing to ciRS-7 acceptor).

### ENCODE data analysis

For ChIP-seq analysis of histone modifications, processed narrowPeak files aligned to hg19 were downloaded from the ENCODE portal. All samples for ChIP-seq were selected with the following filtering criteria, based on annotations in the metadata annotation file downloaded from https://www.encodeproject.org/metadata/type=Experiment&replicates.library.biosample.donor.organism.scientificname=Homo+sapiens/metadata.tsv: “Assay” ⩵ “ChIP-seq”, “File format” ⩵ “bed narrowPeak”, “Output type” ⩵ “peaks”. Only samples with availability of H3K4me1, H3K4me3, H3K9me3, H3K27ac, H3K27me3, and H3K36me3 were selected. For RNA-seq analysis, raw reads were similarly downloaded from the ENCODE portal. Total RNA-seq experimental data were filtered based on the following criteria: “File format” ⩵ “fastq”, “Output type” ⩵ “reads”, “Biosample treatment” ⩵ null, “Library depleted in” ⩵ “rRNA”, “Biosample subcellular fraction term name” ⩵ null. The lists of cell types for which satisfactory ChIP-seq and RNA-seq data were cross-referenced to identify the list of 34 cell types for which data was analyzed.

ChIP-seq narrowPeak file were processed according to the following pipeline: the enrichment scores of any peak calls in genomic 500bp bins spanning 50 kb up- and down-stream of the genomic locus were identified using bedtools intersect -wao -a [genomic_bins.bed] -b [narrowPeak file]. If multiple peaks were called in a given bin, the peak with the highest enrichment score was reported using bedtools merge -c [enrichment column] -o max. The absence of a peak was reported as a 0, which could indicate either complete lack of signal or insufficient signal over input to call a significant peak in the ENCODE pipeline. Fold enrichment values for replicate ChIP-seq experiments performed on the same cell type were averaged for a given genomic bin.

RNA-seq reads were quantified using kallisto quant against a custom kallisto index consisting of RefSeq cDNA transcripts, exclusive of any covering the ASINC/CiRS-7 locus, plus sequences corresponding to individual ASINC/ciRS-7 exons transcribed from both the plus and minus strands[28]. For single-ended data, the quant --[fr/rf]-stranded -b 0 -t 2 -l 200 -s 20 --single command was used, using either --fr-stranded or --rf-stranded depending on the order of the input files. For paired-ended data, the quant --[fr/rf]-stranded -b 0 -t 2 command was used, again using either --fr-stranded or --rf-stranded depending on the input read files.

Expression data was quantified as 1000*tpm/transcript length. Expression values (RPKM) were averaged across replicates for a given cell type for a given transcript.

Our methodology for identifying novel promoters is described below and fundamentally differs from typical informatic approaches such as machine-learning algorithms that use hidden variables or neural networks. These approaches have unknown statistical properties such as effective degrees of freedom or an easily-modeled null distribution for the final test statistic. Below, we describe a method that is conceptually simple and statistically transparent in that the null distribution can be easily computed, and our statistic of interest, numerically related to the active promoter, can be referred to this distribution to obtain an empirical p value.

Decoy ChIP marks on chromosomes 7, 12, and X, corresponding to regions +/− 50 kb upstream and downstream of the annotated TSS and transcription stop sites respectively in the ACTB, FOXO4, and HOTAIR loci, were used as an empirical null for determining the false-discovery rate (FDR) for correlations between ChIP enrichment in putative promoters and enhancers (S3 File). We computed an FDR as follows. For each mark, we computed the correlation between ciRS-7, measured as exonic TPM, and chromatin mark enrichment for bins on chromosomes 7, 12, and X, and generated the empirical distribution of these correlations (S5 File). Then for each mark, we determined the correlation value above which 0.5% of the data in the empirical null fell (q<0.005), and assigned a correlational threshold on the basis of this. We model-checked our assumptions for the empirical null distribution by showing that there was no significant correlation or anticorrelation between ciRS-7 and ACTB (Pearson r = 0.31, p-value = 0.08), ciRS-7 and HOTAIR (Pearson r = −0.13, p-value = 0.45), or ciRS-7 and FOXO4 (Pearson r=0.12, p-value 0.48) as such effects could distort our null model. Heatmaps were generated by averaging ChIP-seq peak enrichment for a given 500bp genomic bin across all sample replicates, and computing the Pearson correlation coefficient of ChIP enrichment against RNA-seq expression for a given cell type.

For genome browser screenshots, pre-processed data was obtained from the ENCODE portal (see S6 File) and visualized using the UCSC genome browser. Data were shown as the mean value over a smoothing window of 6 pixels.

### Quantification of novel junctions and isoforms from ENCODE RNA-seq data sets

370 paired-end RNA-sequencing data sets made from ribosomal RNA-depleted total RNA were downloaded from the ENCODE portal. This set included some samples for which there were biological or technical replicates, representing 136 different cell/tissue types. FASTQ files were quantified against a kallisto index made from ENSEMBL release 89 ncRNA and cDNA fasta files using default parameters (kallisto index -k 31). A second custom kallisto index was created from 40bp junctional sequences within the ASINC locus containing 20bp sequence upstream of a splice site and 20bp sequence downstream of a splice site. This index was also created using default parameters. Paired end data was downloaded and processed using kallisto quant -b 0 against both indexes. tpm (transcripts per million) outputs from the reference transcriptome were aggregated, and for samples with multiple sequencing data sets, tpms were averaged across replicates. For gene-level analysis of the ASINC locus, the five transcripts corresponding to ‘LINC00632’ in ENSEMBL release 89 were summed to determine gene-level expression of the locus. For the custom junctional index, est_counts output for each sample from kallisto quant were normalized to sequencing depth calculated from the reference transcriptome and averaged across replicates. Only junctions with reads supported by at least one sample were reported.

### Analysis of mouse ciRS-7 knock-out data

Data from Piwecka et al. [10] were downloaded from the NCBI Sequence Read Archive using the tool fastq-dump. Data were quantified using kallisto quant --single -l 200 -s 30 against a kallisto index built using default parameters. The kallisto index was generated from a concatenation of cDNA and ncRNA from ENSEMBL release 90 with the addition of the C030023E24Rik sequence, which was not present in the ENSEMBL fasta file. Kallisto quantification was imported into R using tximport [29] and differential gene expression analysis was performed using DESeq2 with default parameters [30]. Aggregation of transcript annotations to perform gene-level analysis was performed with the tx2gene parameter of tximport based on transcript-gene pairing information parsed from the ENSEMBL fasta files. Unless otherwise noted, cutoffs for significance were based genes having an adjusted p-value (“padj”) lower than 0.05.

### Bacterial Artificial Chromosomes (BACs) and plasmid vectors

BACs were purchased from Thermo Fisher Scientific (Waltham, MA) in the case of BAC CTD-2166E9, and from the BACPAC Resources Center (Children’s Hospital Oakland Research Institute, CA) in the case of all other BACs. BACs were purified from E. coli using the Nucleobond Xtra BAC Maxi Kit (Macherey-Nagel, Duren, Germany). SP-dCAS9-VPR (Addgene ID: 63798) was provided by the Qi lab[19], and the sgRNA-encoding plasmid along with the Cas9 plasmid, pMCB306 (Addgene ID: 89360) and lentiCas9-Blast (Addgene ID: 52962) respectively, were gifts from Michael Bassik [31,32]. Guides were cloned into pMCB306 cloned into the BlpI/BstXI site using annealed oligos with the appropriate sticky ends (S1 File).

### BAC Fingerprinting

To ensure BACs had the proper insert, 3 μg of each BAC were digested with 12 units of Ban I (NEB, Ipswich, MA) in the manufacturer’s buffer for 1.5 hours at 37°C. The digests were then heated to 65°C for 20 min. The digestion fragments were separated by loading 750 ng per lane on a 1% LE Agarose (GeneMate) gel with 0.5X TAE running buffer. The DNA was visualized with ethidium bromide staining. Simulated BAC fingerprints were created using SnapGene software (from GSL Biotech) (S16 Fig).

### Transfections

All transfections were performed in 6-well plates using 7.5 μL of Lipofectamine 3000 and 2.5 μg of total DNA per well according to the manufacturer’s protocol. Unless noted otherwise, cells were harvested 24 hours after transfection.

For CRISPRa experiments, we introduced 1.25 μg of the SP-dCAS9-VPR plasmid and 1.25 μg of combined sgRNA plasmids (four for each promoter tested) (see S1 File for sequences). Cells were harvested 48 hours after transfection.

For CRISPR genomic deletions, two sgRNA vectors (pMCB306) with guides targeting a ~55 kb deletion of the X chromosome were transfected into HEK293T cells at a total mass of 1.25 mg per transfection along with 1.25 mg of a vector expressing Cas9 (lentiCas9-Blast). The sequences for these guides can be found in S1 file. The cells were incubated for two days prior to being sorted into single cells by GFP fluorescence as a measure of transfection, which is also expressed on pMCB306. After two weeks, colonies were screened by PCR for the presence of the deletion.

### Nuclear/Cytoplasmic Fractionation

1-2×10^6^ 2 93T cells were fractionated for nuclear/cytoplasmic RNA using the PARIS kit (Thermo Fisher Scientific) according to the manufacturer’s instructions. 0.25-0.5 μg of RNA of each fractionated sample was used in the RT prior to qPCR, using an equal RNA mass for both the nuclear and cytoplasmic fractions in each experiment.

### RNA Purification (primary tissue)

Snap-frozen total brain tissue from a 12-week old C57BL/6 pregnant female mouse was homogenized in TRIZOL and the aqueous phase was purified on Purelink RNA column with on-column DNase treatment.

### RNA Purification (cell lines)

Cells were lysed directly in tissue culture plates by the direct addition of TRIzol reagent (Thermo Fisher Scientific). The manufacturer’s protocol was followed with the following modifications: after isolation of the aqueous phase, 1 volume of 100% EtOH was added to the sample, and then the entire volume was applied to and spun through a RNA Clean & Concentrator-5 column (Zymo, Irvine, CA). The column protocol was performed as per the manual’s instructions starting from the application of the RNA Prep Buffer.

### RNase R treatment

1 μg of RNA was treated with 5 U RNase R (Epicentre, Madison, WI) (or no enzyme in the case of the mock) in 10 μL total reaction volume at 37 C for 30 min. 1 μL 1 mM EDTA, 10 mM dNTPs, and 25 uM random hexamers were then added to the sample and the sample was then heated to 65 C for 5 min to denature RNA structures. The RNA was then reverse transcribed without purification with the addition of 4 μL 5x supplement buffer (250 mM Tris pH 8, 125 mM KCl, 15 mM MgCl2) and 2 μL of 0.1 M DTT to provide the necessary conditions for the RT reaction.

### RT-PCR and qPCR

Total RNA was reverse transcribed with random hexamers using 100 U Maxima Reverse Transcriptase (Thermo Fisher Scientific) according to the manufacturer’s instructions. Endpoint PCRs were performed using DreamTaq DNA Polymerase (Thermo Fisher Scientific, Waltham, MA). For all PCR reactions, 1.5 μl of the unpurified RT-reaction was used per 50 μl reaction volume. All RT-PCR reactions were performed using the recommended cycling protocol for 35 cycles.

qPCR reactions were assembled as 10 μl reactions using AccuPower 2X GreenStar qPCR Master Mix (Bioneer, Daejeon, Korea) with 0.3 μl of template used per reaction. qPCRs were performed on an ABI 7900HT using following cycling protocol: 50 °C for 20 min, 95 °C for 10 min, (95 °C for 15 s and 60 ° C for 60 s) × 45 cycles, followed by a dissociation stage.

### Northern Blot

Northern Blots were performed using 5 μg of total RNA per well using the NorthernMax kit (Thermo Fisher Scientific) according to the manufacturer’s recommendations. Single-stranded DNA oligos were used as probes and were purchased from IDT (Coralville, IA). Probe sequences can be found in S1 File.

## Supporting Information

**S1 Fig. Additional ChIP-seq plots across the ciRS-7 locus from neuronal and HeLa cells.**

**S2 Fig. Heatmap of pearson correlations between chip marks at null loci and ciRS-7 expression:** ACTB (top), HOTAIR (middle), FOXO4 (bottom). Coordinates reported are from the hg19 genome build.

**S3 Fig. Sample correlational plots of chip mark enrichment vs ciRS-7 expression at the distal promoter region.**

**S4 Fig. CRISPR activation of LINC00632 promoters induces the expression of ciRS-7 in HeLa cells.** qPCR measurement of ciRS-7 expression relative to actin. Error bars represent the standard deviation of biological replicates.

**S5 Fig. RNase R sensitivity of transcripts generated from BAC O vs HEK293T.** LINC00632 isoform T3 was measured in both cases. Error bars represent the standard deviation of biological replicates.

**S6 Fig. Transfection efficiency of the BACs** relative to BAC O quantified by DNA-qPCR of BAC backbone DNA from HeLa cells transfected with the BAC. Error bars represent the standard deviation of biological replicates (with error propagated from BAC O).

**S7 Fig. qPCR quantification of ciRS-7 and LINC00632 isoform T3 in HeLa transfected with BACs and HEK mutants.** (A) RNA expression in HeLa transfected with BACs A, B, and C. All values have been normalized to those for BAC A, and error bars represent the standard deviation of biological replicates (with error propagated from BAC A). (B) RNA expression of isoforms in wild-type HEK293T and in a cloned strain of HEK293T in which the putative ciRS-7 promoters have been deleted.

**S8 Fig. Additional PCRs to determine connectivity between exons of LINC00632 and ciRS-7.**

**S9 Fig. Sanger sequencing traces for novel linear ciRS-7 junctions in human.**

**S10 Fig. Products from TOPO cloning** of bands in Fig 2B--*left* (lanes 2 and 3).

**S11 Fig. qPCR ΔCt of human isoforms.** Higher values indicate lower expression. Error bars represent standard deviation of biological replicates.

**S12 Fig. RNase R sensitivity of ciRS-7, LINC00632, and LINC00632-CDR1AS isoforms in HEK293T.**

**S13 Fig. Quantification of ciRS-7 and LINC00632 transcripts across ENCODE tissues and cell lines.** (A) LINC00632 and ciRS-7 gene-level quantification, transcripts per million reads (tpm) (B) ciRS-7 backsplice and LINC00632-ciRS-7 junctional counts per million reads

**S14 Fig. qPCR ΔCt of mouse isoforms** *(Top*) ΔCt (vs GAPDH) for ciRS-7 linear and circular isoforms and (*Bottom*) RNase R sensitivity of transcripts in mouse brain. Error bars represent standard deviation of technical replicates.

**S15 Fig. Volcano plots of log fold-change ciRS-7 KO vs WT in each brain region (Methods).**

**S16 Fig. BAC quality checks.** (A) Simulated and experimental BanI digest of the four BACs used in this study. The agreement of these footprints supports that BAC inserts are as reported and have not been significantly altered by bacterial recombination. (B) Sanger sequencing of the 5’ ends of BAC genomic inserts. The vector sequence is in lowercase; the genomic insert sequence is in uppercase.

**S1 File. PCR primers, Northern probes, and sgRNA sequences.**

**S2 File. RNA-seq scripts and output for novel isoform discovery.**

**S3 File. Table of ChIP-seq peak enrichment and RNA expression levels.**

**S4 File. Quantification of RNA expression in ENCODE data.**

**S5 File. Calculation of empirical FDR for ChIP-seq and RNA-seq correlations.**

**S6 File. ENCODE accession numbers used in genome browser screenshots.**

## Acknowledgments

We thank Peter Wang, Peter Sarnow, Zoe Davis and Robert Bierman for discussion and critical comments that improved our work; Roozbeh Dehghannasiri for help with analysis, the Krasnow and Herschlag Labs for discussions, sharing reagents and space; the Qi and Bassik labs for plasmids, and Inga Jarmoskaite for her help with radiation safety training.

